# Molecular and metabolomic characterization of hiPSC-derived cardiac fibroblasts transitioning to myofibroblasts

**DOI:** 10.1101/2023.10.08.561455

**Authors:** Raghu S Nagalingam, Farah Jayousi, Homa Hamledari, Dina H Baygi, Saif Dababneh, Chloe Lindsay, Ramon Klein Geltink, Philipp F Lange, Ian MC Dixon, Robert A Rose, Michael P Czubryt, Glen F Tibbits

## Abstract

1.

Mechanical stress and pathological signaling trigger the activation of fibroblasts to myofibroblasts, which impacts extracellular matrixcomposition, disrupts normal wound healing,andcan generate deleterious fibrosis (Bohl et al., 2008; Sutton and Sharpe, 2000). Myocardial fibrosis independently promotes cardiac arrhythmias, sudden cardiac arrest, and contributes to the severity of heart failure (Frangogiannis, 2021). Fibrosis can also alter cell-to-cell communication and increase myocardial stiffness which eventually may lead to lusitropic and inotropic cardiac dysfunction (PMID: 33135058). Human induced pluripotent stem cell derived cardiac fibroblasts (hiPSC-CFs) have the potential to enhance clinical relevance in precision disease modeling by facilitating the study of patient-specific phenotypes. However, it is unclear whether hiPSC-CFs can be activated to become myofibroblasts akin to primary cells, and the key signaling mechanisms in this process remain unidentified. We hypothesize that the passaging of hiPSC-CFs, like primary cardiac fibroblasts, induces specific genes required for myofibroblast activation and increased mitochondrial metabolism. Passaging of hiPSC-CFs from passage 0 to 3 (P0 to P3) and treatment of P0 with TGFβ1 was associated with a gradual induction of genes to initiate the activation of these cells to myofibroblasts, including collagen, periostin, fibronectin, and collagen fiber processing enzymes with concomitant downregulation of cellular proliferation markers. Most importantly, canonical TGFβ1 and Hippo signaling component genes including TAZ were influenced by passaging hiPSC-CFs. Seahorse assay revealed that passaging and TGFβ1 treatment increased mitochondrial respiration, consistent with fibroblast activation requiring increased energy production, whereas treatment with the glutaminolysis inhibitor BPTES completely attenuated this process. Based on these data, the hiPSC-CF passaging enhanced fibroblast activation, activated fibrotic signaling pathways, and enhanced mitochondrial metabolism approximating what has been reported in primary cardiac fibroblasts. Thus, hiPSC-CFs may provide an accurate in vitro preclinical model for the cardiac fibrotic condition, which may facilitate the identification of putative anti-fibrotic therapies, including patient-specific approaches.

**Highlights:**

- Passaging promotes the activation of fibroblasts to myofibroblasts.
- TGFβ1 treatment activates the fibroblasts, but their expression profile was uniquely different from myofibroblasts.
- High energy requiring fibroblast activation is dependent on glutaminase-based mitochondrial metabolism.
- Passaging induces TGFβ1 and Hippo signaling pathways in activated fibroblasts and myofibroblasts.

**Graphical Abstract Caption:** Probing the activation of fibroblasts to myofibroblasts is key in ECM remodeling processes to avoid fibrosis-related adverse complications, and to better understand disease pathology. Here we report that passaging of hiPSC-derived cardiac fibroblasts promotes fibroblast activation along with a gradual shift in gene expression and metabolic changes towards myofibroblasts. TGFβ1 treatment activates non-passaged fibroblasts, but they are dissimilar to myofibroblasts. The energy-intensive fibroblast to myofibroblast activation process is dependent on glutaminase-mediated mitochondrial metabolism and is prevented by treatment with GLS-1 inhibitor BPTES. Our work demonstrates that hiPSC-CFs can offer a preclinical model analogous to primary cardiac fibroblasts that is comparable with passage-mediated myofibroblast activation and increased mitochondrial metabolism. hiPSC-CFs may also facilitate patient-specific novel anti-fibrosis drug screening and disease management.

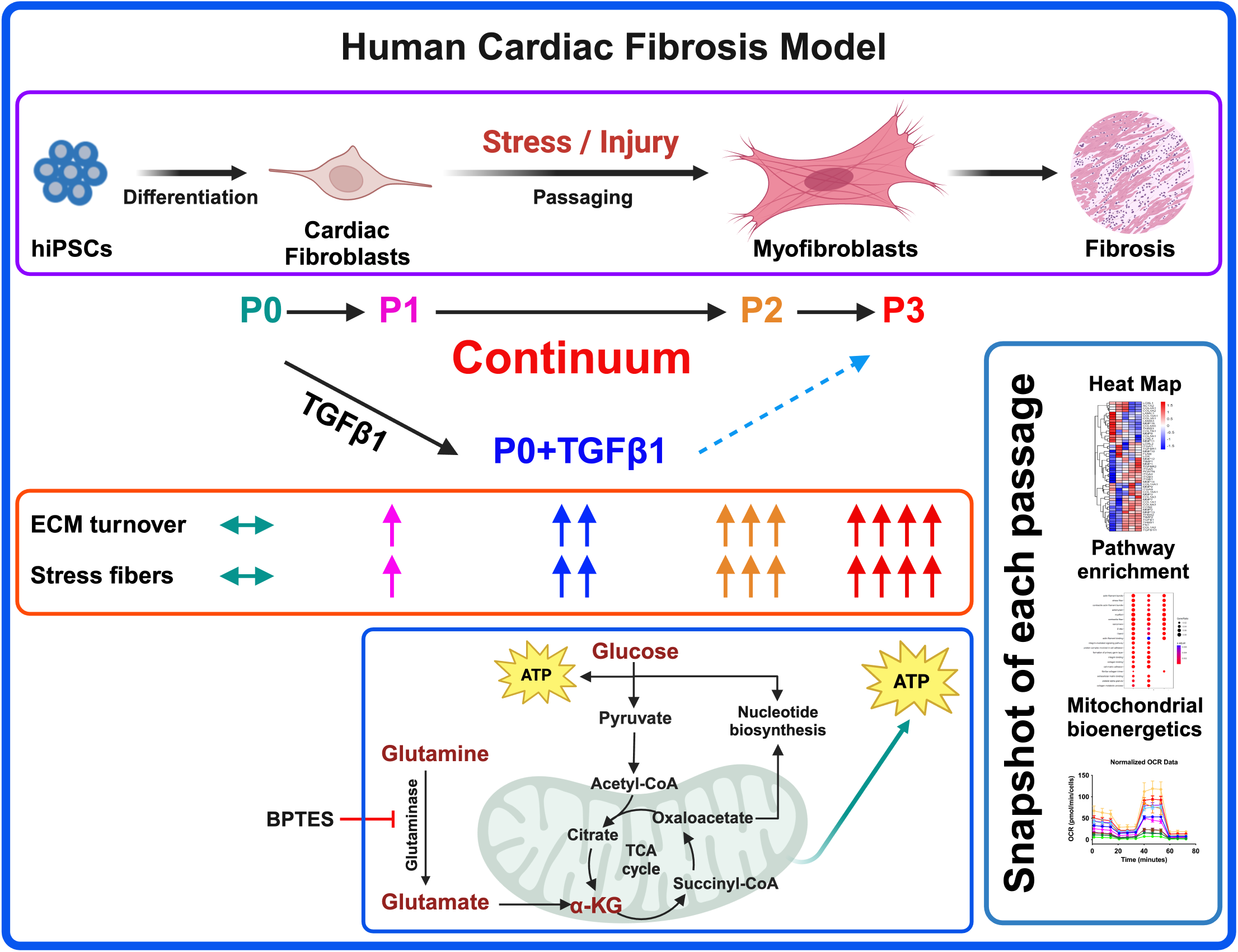

## 2. Introduction

Extracellular matrix (ECM) remodeling in the heart is crucial to forming a stable scar after myocardial infarction (MI) and contributes to replacement fibrosis after a significant loss of cardiomyocytes upon injury (Bohl *et al*., 2008; Sutton and Sharpe, 2000). In hypertensive and diabetic myocardia, uncontrolled ECM alteration can lead to reactive fibrosis upon activation of renin-angiotensin and β-adrenergic signalling (Tanaka et al., 1986; Weber and Brilla, 1991; Weber et al., 1988). ECM remodeling is tightly coordinated, and several trophic factors are known to stimulate cardiac fibroblast activation. Wound healing, in the absence of fibrosis, is marked by fibroblast activation to myofibroblasts (Frangogiannis, 2021) and rapid scar formation followed by a period of myofibroblast apoptosis which results in healed tissue characterized by continuous and relatively slow matrix turn-over. Conversely wound healing after myocardial infarction, or in response to prolonged stress in the heart such as in pressure overload, is typified by the relative persistence of myofibroblasts with attendant excessive synthesis and deposition of ECM and matrix-associated proteins (Hinz and Lagares, 2020; Nagalingam et al., 2022). This excessive deposition includes structural collagens as well as periostin (Shimazaki et al., 2008) and the fibronectin ED-A splice variant. As our understanding of the pathophysiology and signaling mechanisms of cardiac fibroblast activation and the genes that are associated with this activation remains incomplete, its investigation is warranted. Several groups have investigated the signaling mechanisms governing fibroblast activation using primary murine cardiac fibroblasts (Klingberg et al., 2018). The expression of α-smooth muscle actin (α-SMA) by myofibroblasts is often cited as a standard marker protein of fibroblast activation and after acute MI, increased numbers of α-SMA-positive myofibroblasts are found within the infarct region (Virag and Murry, 2003). Lineage tracing of cardiac myofibroblasts in a murine MI model revealed that after scar formation, some myofibroblasts transition towards quiescent matrifibrocytes, and that in the human myocardial scar, similar matrifibrocytes with unique gene expression and secretome exist and are separate from myofibroblasts (Fu et al., 2018). Whether the persistence of myofibroblasts and appearance of matrifibrocytes occurs in hearts of various etiologies of disease and in non-murine preclinical models remains an open question. In a left ventricular pressure overload model, α-SMA expressing myofibroblasts were found to be elevated at 2 weeks and then declined after 4 weeks, and in vitro cultured fibroblasts revealed that α-SMA expression peaked at 9 days and declined by 12 days, indicating that fibroblast activation may present as a dynamic transition (Gilles et al., 2020). Removal of periostin-positive myofibroblasts results in less severity of the disease phenotype by limiting collagen synthesis and scar formation after MI (Kanisicak et al., 2016). TGFβ1 (Algeciras et al., 2021; Dobaczewski et al., 2010; Saadat et al., 2020; Villalobos et al., 2019; Zeglinski et al., 2016), and hippo signaling(Landry et al., 2021) is also elevated when fibroblasts are activated to become myofibroblasts. Fibroblast activation to myofibroblasts utilizes energy via mitochondrial metabolism (Negmadjanov et al., 2015) and is highly dependent on glutaminolysis and its rate-limiting enzyme GLS1 (Chattopadhyaya et al., 2022; Gibb et al., 2022). Most fibroblast activation studies to date were performed either in stable cell lines or murine primary cells due to the scarcity of human tissues to prepare primary cells, and commercially available cardiac fibroblasts are often passaged multiple times prior to sale.

There are multiple protocols published so far to differentiate cardiac fibroblasts from iPSCs; however, most of them share a similar path with various small molecules to initiate the cardiac mesoderm formation followed by epicardial cell conversion to cardiac fibroblasts. To generate the cardiac mesoderm and cardiac progenitors from iPSCs, most protocols used a GSK3 inhibitor (CHIR99021) and a WNT signaling inhibitor (IWR1 or XAV939); after cardiac mesoderm formation, a TGFβ-inhibitor (SB431542) and retinoic acid (RA) promoted conversion to epicardial lineage cells, followed by FGF-2 treatment to generate hiPSC-CFs (Beauchamp et al., 2020; Campostrini et al., 2021; Cumberland et al., 2023; Giacomelli et al., 2020; Hall et al., 2023; Soussi et al., 2023; Whitehead et al., 2022; Yu et al., 2022; Zhang et al., 2022; Zhang et al., 2019a). The differentiation protocols for iPSC-derived lung and dermal fibroblasts differ in the growth factors and chemical signals used on specific lineages to become tissue-specific fibroblasts (Alvarez-Palomo et al., 2020; Itoh et al., 2013; Kim et al., 2018; Mitchell et al., 2023; Tamai et al., 2022; Wong et al., 2015). Most notably, NKX2.5 defines the cardiac-specific lineage from cardiac mesoderm (Zhang *et al*., 2022; Zhang *et al*., 2019a), whereas NKX2.1 directs lung-specific lineage (Mitchell *et al*., 2023; Wong et al., 2012). iPSC-CFs express cardiac-specific genes like GATA4, BMP4, TCF21, DDR2, TE-7, HAND2, HEY1, ISL1 and α-SMA and are influenced by their epicardial lineage (Campostrini *et al*., 2021; Giacomelli *et al*., 2020; Zhang *et al*., 2022; Zhang *et al*., 2019a; Zhang et al., 2019b). iPSC-derived lung fibroblasts express different sets of genes like NKX2.1, GATA6, DNP63a, FOXA1, ACE1, AQP5, T1α, SPA, SPB, SPC FOXF1, and FOXA2 (Alvarez-Palomo *et al*., 2020; Mitchell *et al*., 2023; Tamai *et al*., 2022; Wong *et al*., 2015); whereas iPSC-derived dermal fibroblasts and keratinocytes presented with not only KRT14, also ΔNp63, DSG3, ITGB4, laminin 5, KRT5, KRT1 and loricrin (Itoh *et al*., 2013; Kim *et al*., 2018).

Current advancements and optimization in protocols that are more efficient in generating hiPSC-derived cardiac fibroblasts provide a compelling model to mimic myofibroblast activation to study fibrosis. A recent study reported the use of TGFβ1 inhibitors to maintain the fibroblasts as quiescent for up to five passages (Zhang *et al*., 2019a). Fibroblasts locally generate TGFβ1 during mechanical stretch upon interacting with integrins (Munger et al., 1999; Sarrazy et al., 2014) and TGFβ1 is a critical player in activating the fibroblasts and fibrosis-associated ECM gene expression in cultured fibroblasts (Dobaczewski *et al*., 2010; Eghbali et al., 1991; Heimer et al., 1995; Sarrazy *et al*., 2014; Villarreal et al., 1996; Yi et al., 2014). Several studies used passaging and TGFβ1 to induce fibroblast activation and the myofibroblast phenotype. It is unclear whether TGFβ1-induced fibroblast activation is similar to the myofibroblastic phenotype produced by passaging. In this study, our goal was to capture a snapshot of passage-mediated cardiac fibroblast to myofibroblast transition and TGFβ1-induced fibroblast activation in hiPSC-derived cardiac fibroblasts using transcriptomic, proteomic, and metabolomic profiling and signaling mechanisms involved to provide a better understanding of the fibrosis-associated disease phenotype.

## 3. Results

### 3.1. Passaging promotes fibroblast activation to myofibroblasts

Activation of fibroblasts to myofibroblasts is crucial for normal wound healing and elevated ECM turnover for stable scar formation after MI (Frangogiannis, 2021). In our hiPSC-derived cardiac fibroblasts, upon passaging towards the myofibroblast (P3) phenotype or treatment with TGFβ1 to become activated fibroblasts (P0+TGFβ1), both phenotypes display a reduction in the expression of the proliferation markers CDK-4 at the mRNA level (**Fig. S1-A**) and CDK-1, MCM3, and PCNA at the protein level (**Fig. S1-B, C, D**) compared to the non-passaged fibroblasts (P0). The characteristic hallmark of fibroblast to myofibroblast activation is α-SMA positive stress fiber incorporation inside the myofibroblasts (Hinz, 2007). Phalloidin staining and DDR immunofluorescence revealed that hiPSC-derived cardiac fibroblasts that underwent three passages (P1 to P3) or P0+TGFβ1 showed gradual expression of stress fibers in higher passaged cells compared to non-passaged cells (P0) (**Fig. 1-B**). Most importantly, after passaging hiPSC-CFs displayed a reduction in TCF21 gene expression from P0 to P3 and in P0+TGFβ1, whereas myofibroblast marker genes (e.g., periostin (POSTN) and ED-A-fibronectin (ED-A-Fn)) showed an incremental expression pattern in P0 to P3 and in P0+TGFβ1 compared to P0 cells at the mRNA level using qPCR analysis (**Fig. 1-C-E**). These results clearly indicate that hiPSC-derived fibroblasts, upon passaging, gradually become myofibroblasts.

**Figure 1:**
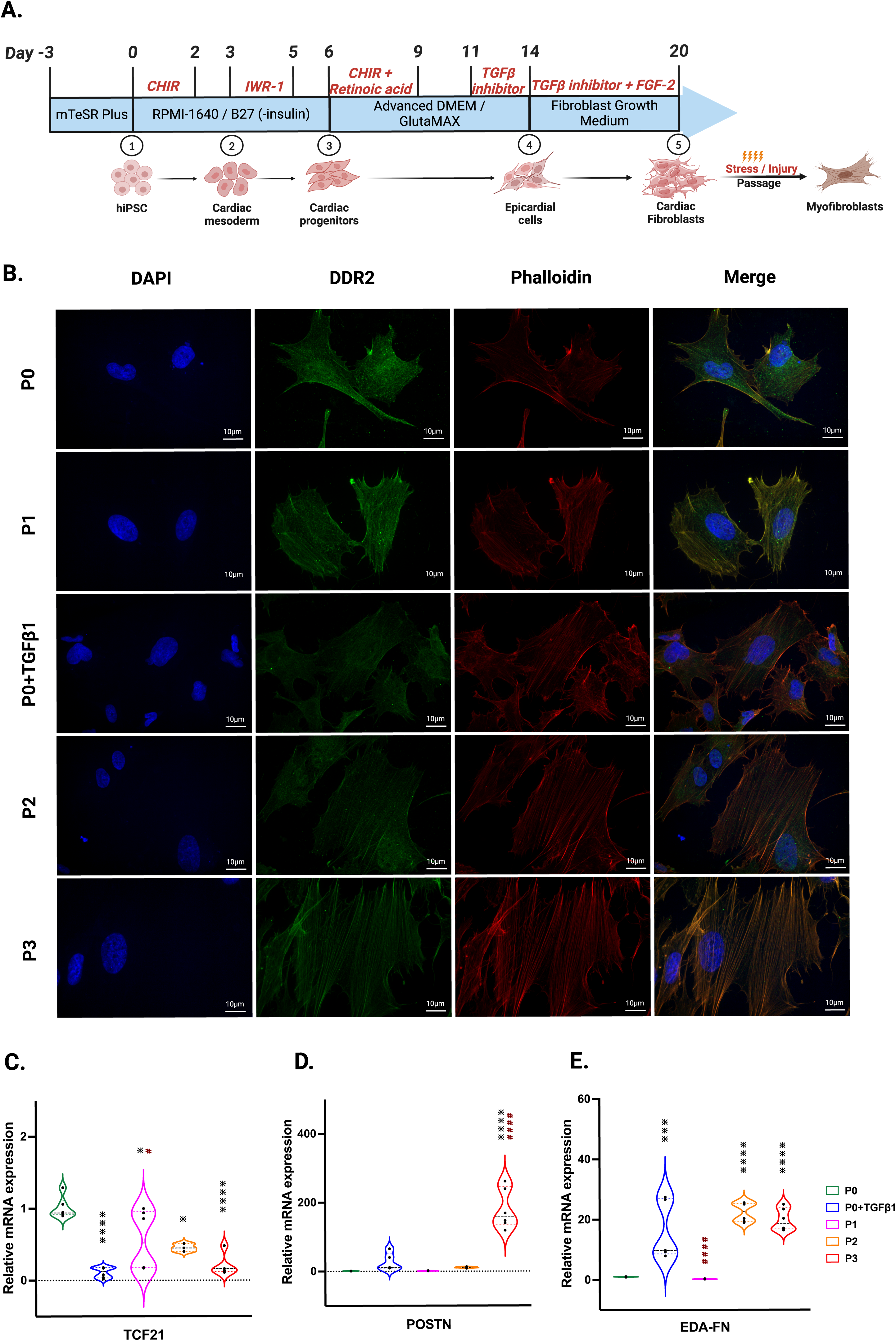
hiPSC-derived fibroblast transition to the myofibroblast phenotype after passaging. Timeline and differentiation of fibroblasts from hiPSCs (**A**). Images represent stress fibers stained with phalloidin (red), DDR-2 (green) and DAPI (blue) in hiPSC-derived non-passaged fibroblasts (P0), non-passaged fibroblasts treated with TGFβ (P0+TGFβ1) and different passages of fibroblasts (P1, P2, P3) (**B**). Passaging of fibroblasts initiates myofibroblast activation, altering the mRNA expression of TCF21 (**C**), POSTN (**D**), and EDA-Fn (**E**) as assessed by qPCR. Statistical significance was determined by one-way ANOVA with Tukey’s post-hoc test (**n=3-4**). *p<0.05, **p<0.01, ***p<0.001, ****p<0.0001 vs P0 and #p ≤ 0.05, ####p ≤ 0.0001 vs P0+TGFβ1.

### 3.2. Passaging elevates fibrosis-associated genes while transitioning to the myofibroblast phenotype

We compared the different passages of hiPSC-CFs with the NanoString codeset using a PCA plot displaying unique clusters of various passages of hiPSC-CFs (P0, P1, P2, P3) and TGFβ1 treated non-passaged cells (P0+TGFβ1). The differences are 55.6% and 14.6% in component-1 (X-axis) and component-2 (Y-axis), respectively. The P0 and P3 clusters showed the greatest differences between each other, while other clusters (P1, P2, P0+TGFβ1) fell between P0 and P3 (**Fig. 2-A**). The group correlation of gene expression between various passages of hiPSC-CFs were plotted based on Pearson correlation, and the higher intensity of red corresponds to high correlation, and higher green corresponds less correlation between groups (**Fig. 2-B**). Volcano plots were generated based on fold change between P0+TGFβ1 vs. P0 (**Fig. 2-C**), P1 vs. P0 (**Fig. 2-D**), P2 vs. P0 (**Fig. 2-E**), P3 vs. P0 (**Fig. 2-F**), P3 vs. P0+TGFβ1 (**Fig. 2-G**), in which upregulated genes are depicted on the left side and downregulated genes are on the right side of the panel. Volcano plot analysis revealed that each passage of hiPSC-CFs (P1, P2, P3) and P0+TGFβ1 gene expression pattern is distinct when compared to P0; also, P3 displayed significantly different gene expression compared to P0+TGFβ1. The P3 group showed significant upregulation in profibrotic and myofibroblast-related genes, whereas P1, P2, and P0+TGFβ1 showed a steady shift towards myofibroblast-related profibrotic genes such as POSTN, ED-A-Fn and key fibrillar collagens (Col1α1 and Col1α2), whereas α-smooth muscle actin (ACTA2) expression was higher in the P1 and P0+ TGFβ1 compared to P0 but not in P2 and P3. We tested the differential expression of selective ECM-related genes comparing various passages of hiPSC-CFs (P0, P1, P2, P3) and P0+TGFβ1 using heatmap analysis (**Fig. 2-H**). The heatmap data revealed that the expression profile of different passages of hiPSC-CFs (P1, P2, P3) and P0+TGFβ1 caused a significant shifting towards the myofibroblast phenotype upon passage-mediated fibroblast activation. Activated fibroblasts and myofibroblasts are responsible for the increased ECM turnover in reparative and reactive fibrosis, increasing collagen, fibronectin, and other secretory proteins. Passaging of hiPSC-CFs revealed that gene expression of major fibrotic fibrillar collagens, Col1α1 and Col1α2, was significantly higher in both P0+TGFβ1and P3 compared to P0, whereas Col3α1 showed a reduction in P0+TGFβ1 and P3 compared to P0 (**Fig. S3-A, B, C**). Similarly, gene expression of structural and secretory proteins like α-smooth muscle actin (ACTA2), periostin (POSTN), fibronectin (FN1), vimentin (VIM), integrin beta-1 (ITGB1), and integrin alpha-4 (ITGA4) were significantly elevated in P0+TGFβ1 and P3 compared to P0 (**Fig. S3-D, E, F, G, H, I**). ECM remodeling enzymes play a crucial role in the formation and maturation of stable scars, and their expression was elevated in parallel with a significant accumulation of mature collagen fibers in the failing myocardium. Lysyl oxidase (LOX) acts as a catalyst in linking collagen fibers to become mature collagen, and along with other matrix remodeling enzymes such as MMP2 and MMP1 was upregulated in both P3 and P0+TGFβ1 compared to P0 (**Fig. S2-A, B, C**). Comparative cluster enrichment analysis revealed the top 10 upregulated pathways when comparing P3 vs. P0, P0+TGFβ1 vs. P0, or P3 vs. P0+TGFβ1 (**Fig. 2-H**); similar analysis provided the top 10 downregulated pathways in these comparisons (**Fig. 2-I**).

**Figure 2:**
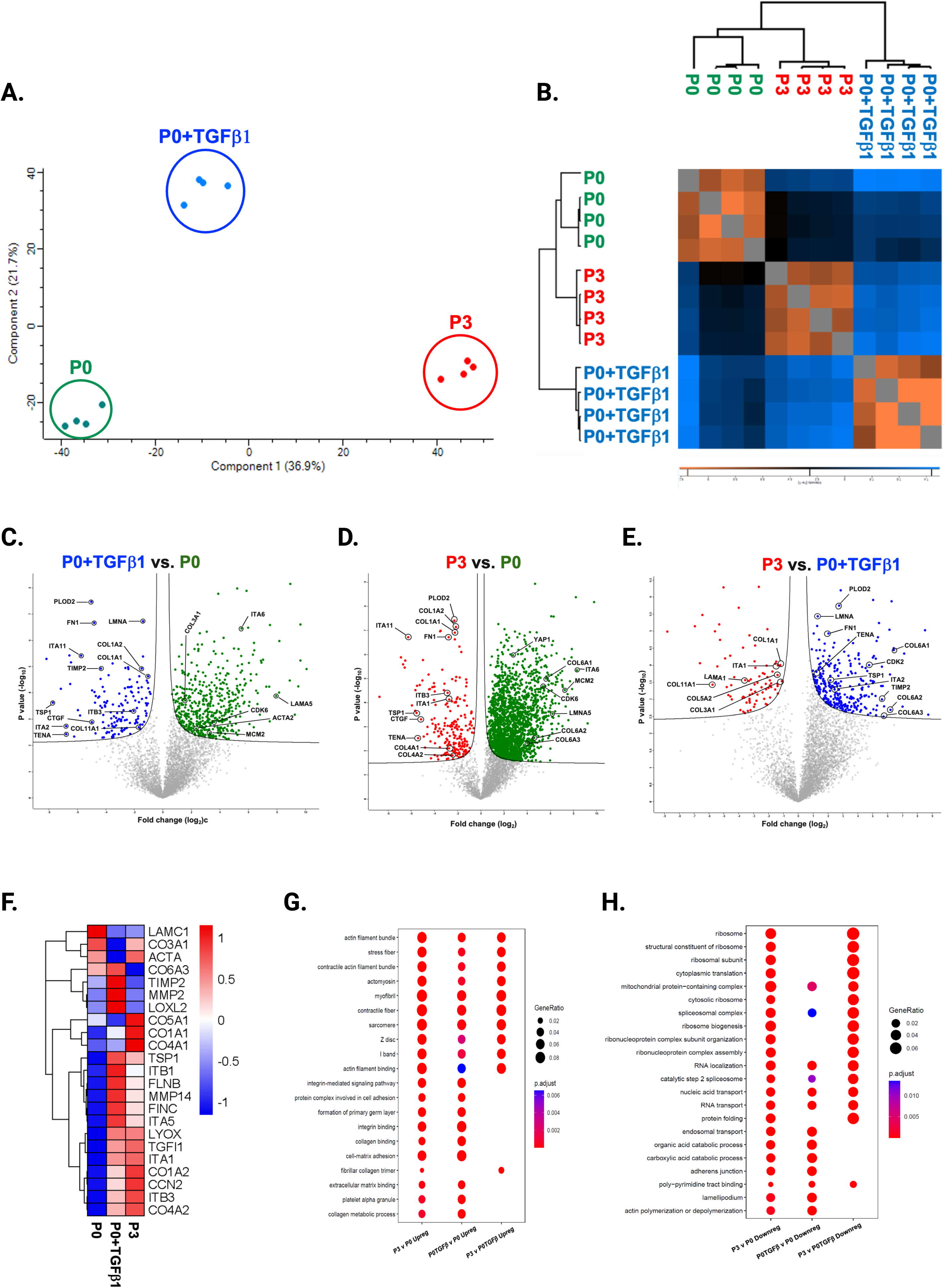
Passaging promotes fibrosis-responsive gene expression. Principle component analysis (PCA) of NanoString data (**A**), based on Benjamini-Hochberg method FDR adjusted to P-value < 0.05. Group correlation of different passages of fibroblasts based on Pearson correlation method (**B**), and the gene expression values were scaled from 86.5 to 98.5. Volcano plot of fibrosis gene expression profile in hiPSC-derived fibroblasts P0+TGFβ1 vs. P0 (**C**), P1 vs. P0 (**D**), P2 vs. P0 (**E**), P3 vs. P0 (**F**), P3 vs. P0+TGFβ1 (**G**), (adjusted P < 0.05), upregulated genes on the left side of the panel and downregulated on the right side (**n=4-5**), per group and the fold-change was compared using Welch’s t-test. Heatmap of differential expression of selected genes comparing various passages of hiPSC-derived fibroblasts (**H**) and the expression values were scaled from -1.5 to 1.5, similarity was calculated using a heatmap package (clustering_distance_rows = "euclidean"). Compare cluster analysis of enrichment of the top 10 upregulated pathways (**I**) and downregulated pathways (**J**) between P3 vs P0, P0+TGFβ1 vs P0, and P3 vs P0+TGFβ1, based on adjusted p-value < 0.05.

### 3.3. Passaging alters the proteomic profile of hiPSC-derived cardiac fibroblasts

To understand the passage-mediated proteome profile changes, we performed principal component analysis (PCA) to see the clustering differences between non-passaged fibroblasts (P0), myofibroblasts (P3), and growth factor activated fibroblasts (P0+TGFβ1) using the Benjamini-Hochberg method (**Fig. 3-A**). The cluster differences are 46.3% in component-1 (X-axis) and 16.9% in component-2 (Y-axis). The P0 and P3 clusters are notably far apart from each other, with P0+TGFβ1 falling in the middle. Group correlation analysis of proteomic expression in P0, P3, and P0+TGFβ1 based on Pearson correlation revealed intensity gradient differences between groups, in which higher intensity of brown corresponds to high correlation and higher blue corresponds minimal correlation (**Fig. 3-B**). Proteome profiles of different passages of hiPSC-CFs (P0, P1, P2, P3) and P0+TGFβ1 were plotted based on fold change between P0+ TGFβ1 vs. P0 (**Fig. 3-C**), P3 vs. P0 (**Fig. 3-D**), P3 vs. P0+ TGFβ1 (**Fig. 3-E**) using volcano plot analysis; these plots indicated that passaging promoted the myofibroblast phenotype in hiPSC-CFs, whereas TGFβ1-treated non-passaged fibroblasts also exhibited distinct proteomic expression compared to either P0 or P3. Differential expression of selected ECM-related proteins compared between P0, P3, and P0+TGFβ1 was examined using heatmap analysis (**Fig. 3-F**), and the data revealed that each group is unique, with expression profiles that differ from one another. Passage-mediated myofibroblasts and TGFβ1-treated hiPSC-CFs revealed that profibrotic proteins such as fibrillar collagens (Col1α1 and Col1α2) and non-fibrillar collagens (Col4α1) were significantly higher in P3 compared to P0, whereas Col3α1 showed a reduction in P0+TGFβ1 and P3 compared to P0 (**Fig. 4-A, B, C, D**). Structural proteins like ACTA2, FN1, integrin α-1 (ITGA1), and integrin β-3 (ITGB3) were significantly elevated in P0+TGFβ1 and P3 compared to P0; however, vimentin (VIM) protein expression went down in P3 compared to P0 (**Fig. 4-E, F, G, H, I**). Additionally, ECM remodeling enzymes such as lysyl oxidase and lysyl hydroxylase-2 (PLOD2) protein expression was upregulated in both P3 and P0+TGFβ1 compared to P0, an indication of elevated collagen processing (**Fig. S2-D, E**). Comparative cluster enrichment analysis of proteomic expression profiles revealed the top 10 up-regulated (**Fig. 3-G**) and down-regulated (**Fig. 3-H**) signalling pathways between (P3 vs. P0), (P0+TGFβ1 vs. P0), and (P3 vs. P0+ TGFβ1); these results indicate that both P3 and P0+TGFβ1 exhibit distinct profibrotic gene expression profiles and signalling when compared to P0.

**Figure 3:**
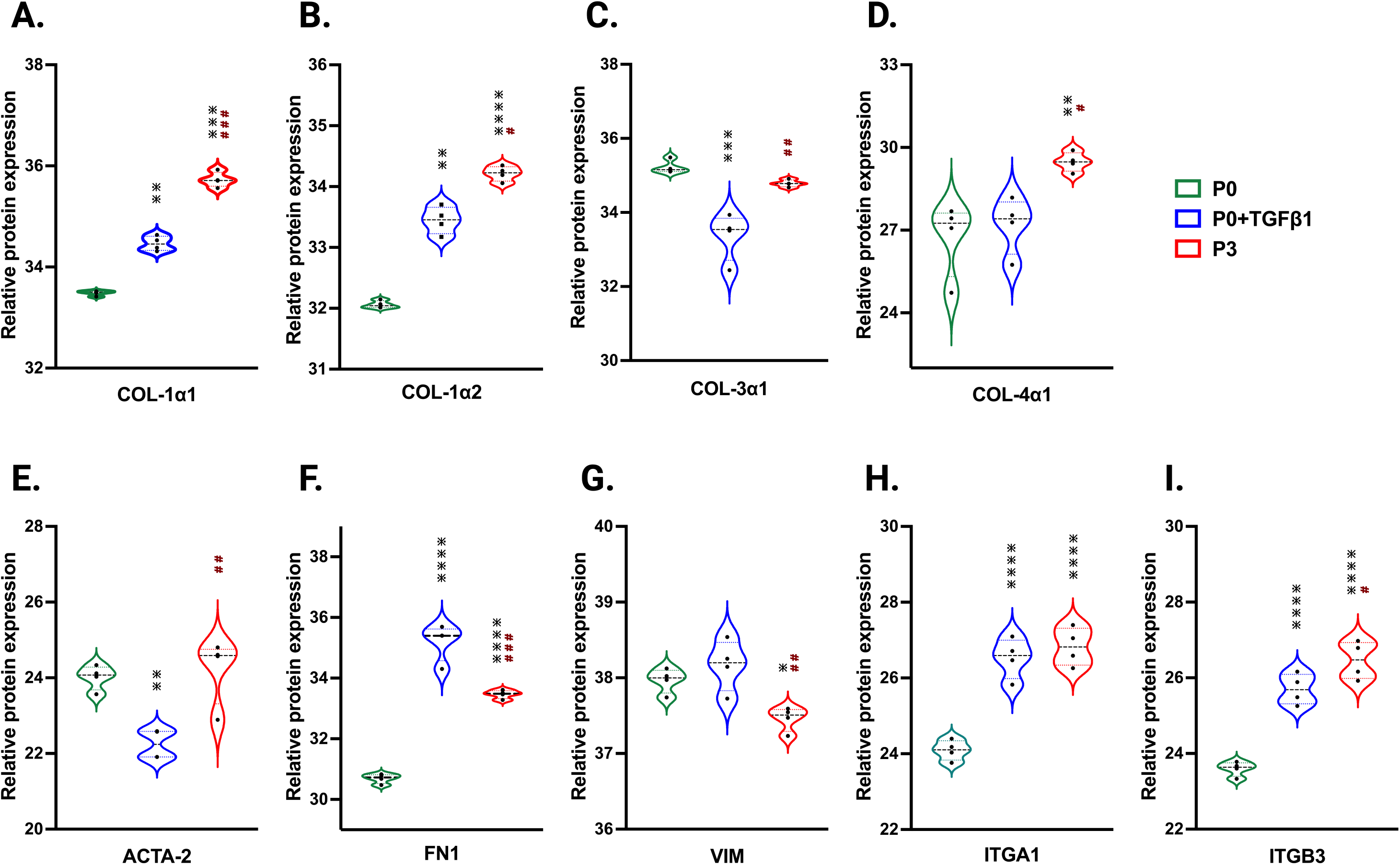
Passaging alters the proteomic expression profile of fibroblasts, activated fibroblasts and myofibroblasts. PCA analysis of proteomic gene expression data by mass spectrometry (**A**), FDR adjusted to P-value < 0.01. Pearson group correlation of different passages of fibroblasts and their gene expression (**B**). Volcano plot of proteomic gene expression profile in P3 vs. P0 (**C**), P0+TGFβ1 vs. P0 (**D**), P3 vs. P0+TGFβ1 (**E**), P value adjusted to < 0.05; (**n=3-4**), per group and the fold-change was measured using Welch’s t-test. Heatmap of differential expression of selective ECM-related genes comparing hiPSC derived P0, P0+TGFβ and P3 (**F**), expression values were scaled from -1.0 to +1.0. Comparative cluster analysis of enrichment of top 10 upregulated pathways (**G**) and downregulated pathways (**H**) between P3 vs P0, P0+TGFβ1 vs P0, and P3 vs P0+TGFβ1, based on adjusted p-value < 0.05.

**Figure 4:**
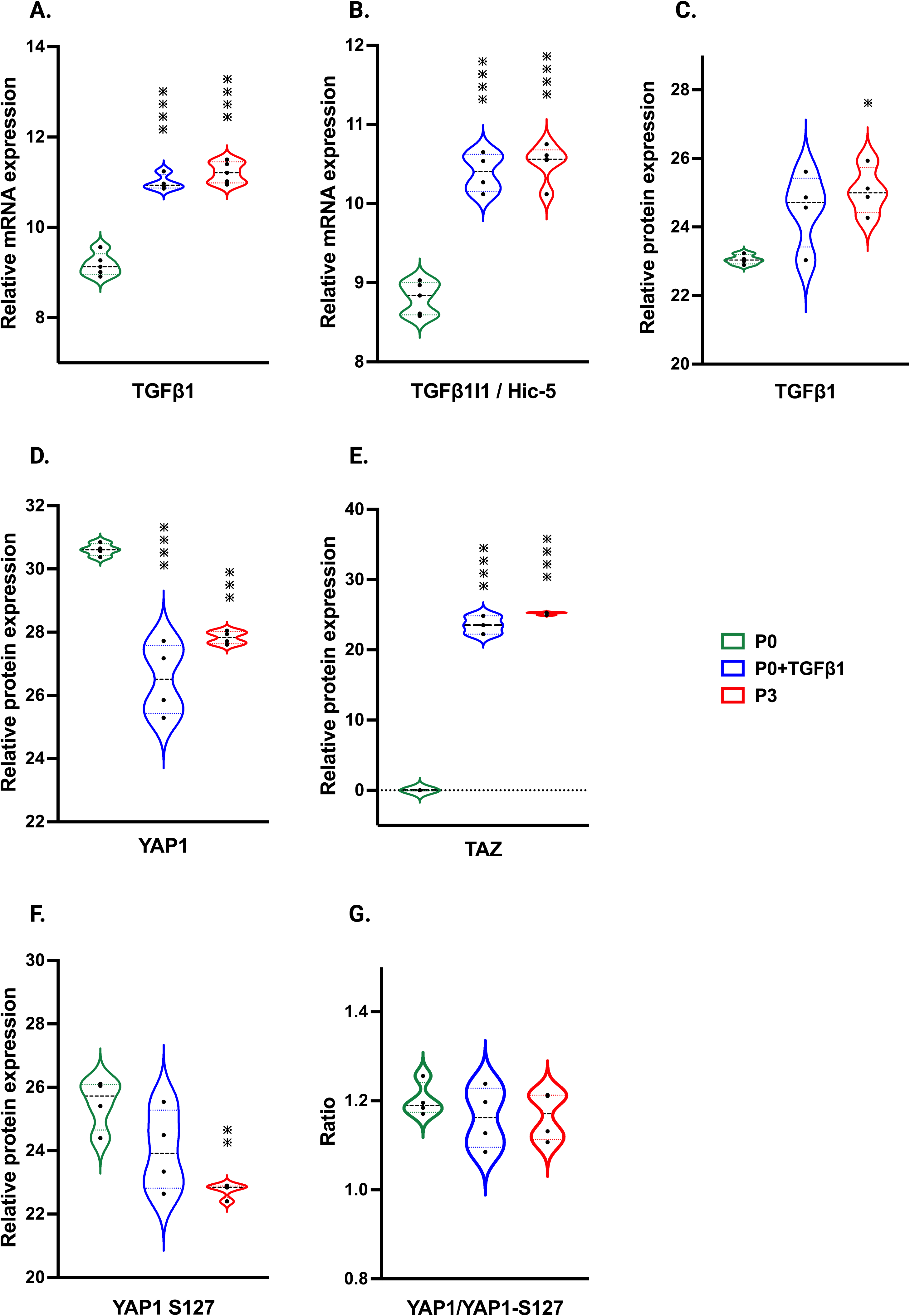
Passaging accelerates the turnover of profibrotic extracellular matrix genes during phenotype. Violin plots of profibrotic ECM protein expression in P0+TGFβ1 and P3 compared to P0, (**A-I**). Statistical significance was determined by one-way ANOVA with Tukey’s post-hoc test (**n=4**); *p<0.05, **p<0.01, ***p<0.001, ***p<0.0001 vs fibroblasts (P0) and #p ≤ 0.05, ##p ≤ 0.01, ###p ≤ 0.001, vs P0+TGFβ1.

### 3.4. Metabolic profiling of hiPSC-derived cardiac fibroblasts transitioning to myofibroblasts

To understand the potential metabolic changes involved in hiPSC-derived cardiac fibroblast to myofibroblast conversion, Seahorse assays for mitochondrial respiration were performed at different passages with/without TGFβ1 treatment. Interestingly, the results of our study showed a significant concomitant increase in both mitochondrial oxygen consumption rate (OCR) (**Fig. 5-A**) and extracellular acidification rate (ECAR) which is a proxy for glycolysis (**Fig. 5-B**) with increased passage number, with the highest values observed in P3 compared to non-passaged fibroblasts (P0). To further understand the role of glutamine metabolism in passage-mediated cardiac myofibroblast phenotype switching in hiPSC-CFs, we treated the various passages of hiPSC-CFs with glutaminase (GLS1) inhibitor BPTES, which limits the conversion of glutamine to glutamate, decreasing energy production via the TCA cycle. We noted a significant attenuation of OCR and ECAR by BPTES, indicating that passage-mediated metabolic changes depended on glutaminolysis, which appears to play an essential role in cardiac fibroblast to myofibroblast transition.

**Figure 5:**
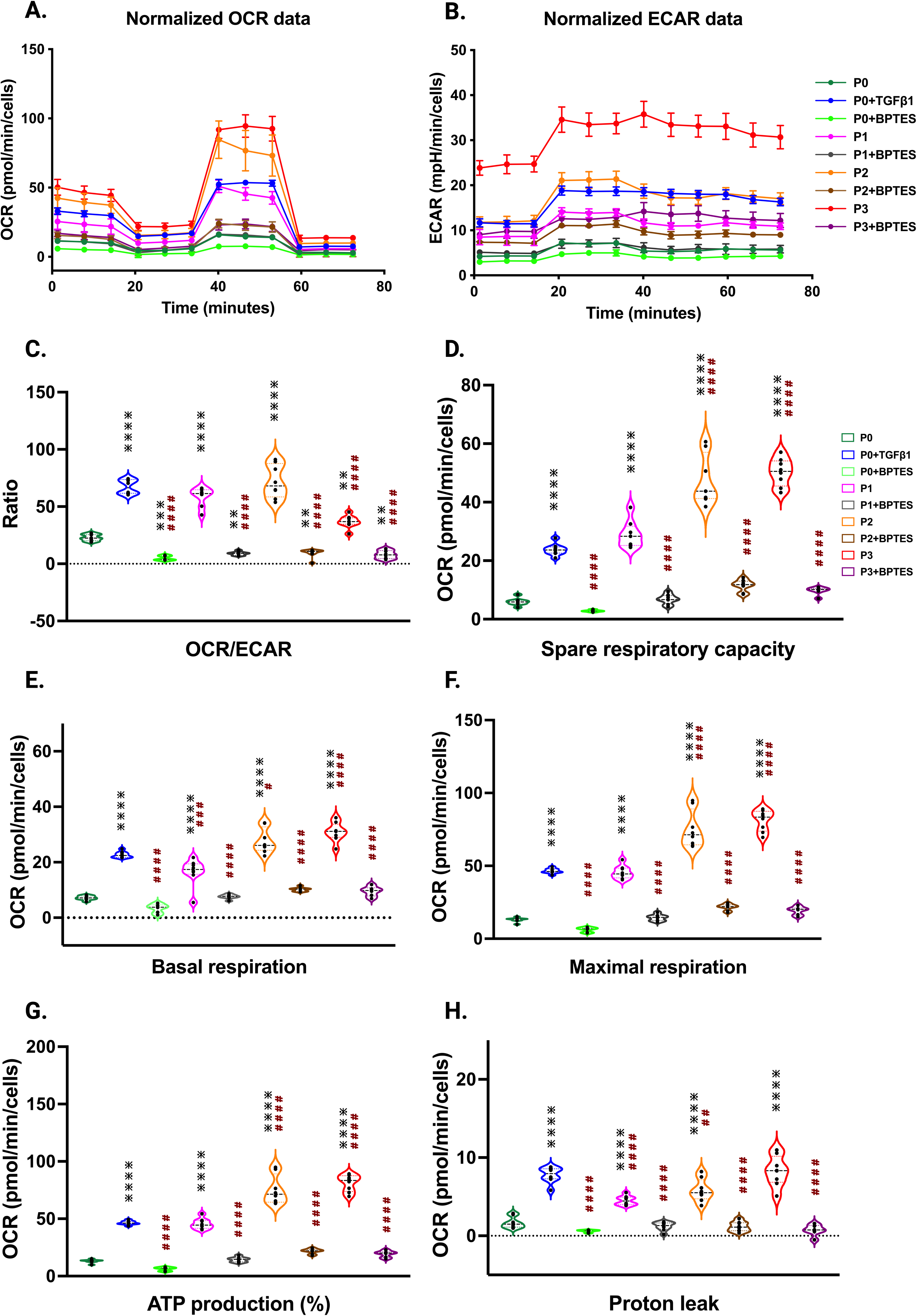
Passage-mediated myofibroblast activation depended on mitochondrial metabolism. Oxygen consumption rate is elevated in higher passage of fibroblasts and in TGFβ1 treatment than no passage fibroblasts Seahorse assay reveals that increased OCR rate upon passage (P3) and/or TGFβ1 treatment, whereas the glutaminase inhibitor BPTES limits this process when compared to non-passaged fibroblasts (P0) (**A-H**), indicating that activation of myofibroblasts primarily depended on the α-keto-glutarate (α-KG) mediated glutaminolysis pathway. Statistical significance for the violin plots was determined by one-way ANOVA with Tukey’s post-hoc test (**n=3-8**); *p ≤ 0.05, **p ≤ 0.01, ***p ≤ 0.001, ****p ≤ 0.0001, vs fibroblasts (P0) and #p ≤ 0.05, ##p ≤ 0.01, ###p ≤ 0.001, ####p ≤ 0.0001 vs P0+TGFβ1.

The OCR/ECAR ratio reveals a more pronounced shift toward mitochondrial respiration than glycolysis for energy metabolism in higher passages with or without TGFβ1, which was inhibited by BPTES treatment (**Fig. 5-C**). The significant boost in spare respiratory capacity in higher passages corresponds to the metabolic adaptability of cells in responding to the increased energy demand during the transition to the myofibroblast stage; however, these responses were reduced with BPTES treatment (**Fig. 5-D**). Similarly, basal and maximal respiration also showed an increased trend toward higher passages with TGFβ1 treatment. ATP production, proton leak, and non-mitochondrial OCR were significantly increased in higher passages and following TGFβ1 treatment compared to P0 (**Fig. 5-E, F, G, H and S4-A**) and these effects were abolished with BPTES treatment. Additionally, we did not observe any differences in coupling efficiency with passage or TGFβ1-mediated fibroblast activation or treatment with glutaminolysis inhibitor BPTES treatment (**Fig. S4-B**).

### 3.5. Signaling mechanisms involved in passage-mediated myofibroblast transition in hiPSC-derived cardiac fibroblasts

In our hiPSC-derived cardiac fibroblasts, upon passage, there was a significant increase in TGFβ1 mRNA and protein expression in both P3 and P0+TGFβ1 (**Fig. 6-A, C**). TGFβ1-induced transcript-1 (TGFβ1I1) also known as Hic-5, involved in activation, stress fiber growth and assembly of myofibroblasts, was highly upregulated in both P3 and P0+TGFβ1 compared to P0 (**Fig. 6-B**). Our hiPSC-derived CFs P0+TGFβ1 and P3 showed a significant reduction in YAP protein expression compared to P0, whereas TAZ expression was upregulated in both P3 and P0+TGFβ1 (**Fig. 6-D, E**). We observed that YAP1 is phosphorylated at S127 site (**Fig. 6-F**), and we also noted no change in the total YAP1 vs YAP1 phosphorylation ratio in P3 and P0+TGFβ1 compared to P0 (**Fig. 6-G**). Phosphorylation of YAP1 promotes the degradation or retains the YAP1/TAZ complex in the cytoplasm to limit the profibrotic signaling. Our activated hiPSC-CFs displayed an increase in the active form of YAP1 than P0; however, we did not observe any change in total YAP1 vs phosphorylated YAP1.

**Figure 6:**
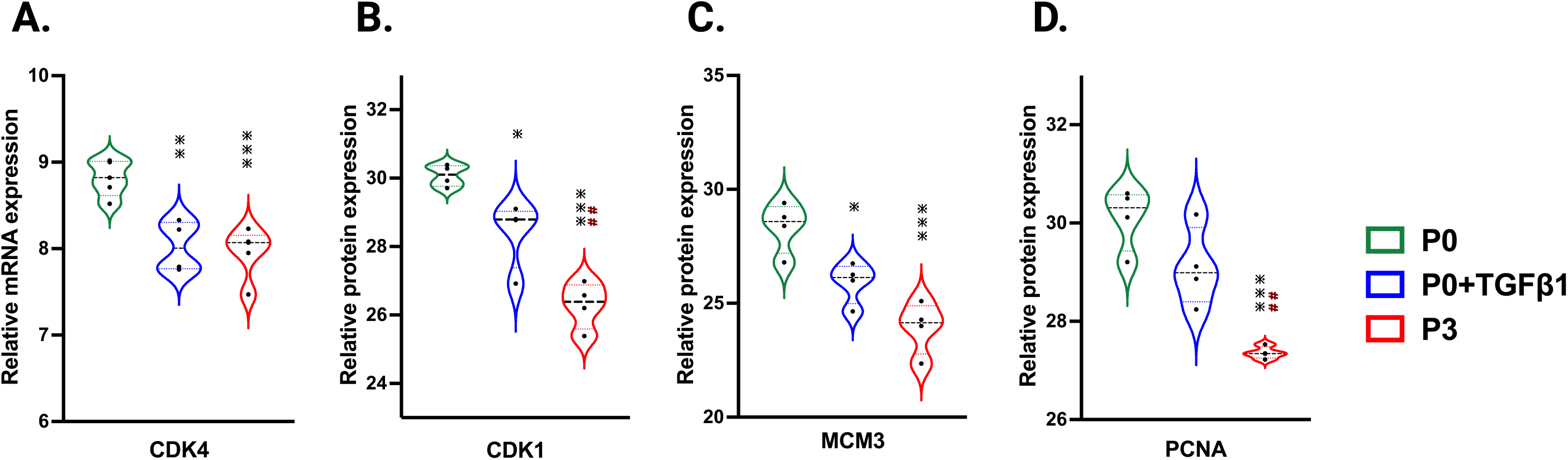
Passaging influences TGFβ1 and Hippo signaling pathway. Violin plots represent mRNA expression of TGFβ1 (**A**), TGFβ1I1 (**B**) and proteomic expression of TGFβ1 (**C**), showing upregulation in the TGFβ1-mediated canonical signaling pathway in P3 and P0+TGFβ1compared to fibroblasts (P0). Proteomic expression of hippo signaling mediators YAP (**D**), TAZ (**E**) and phosphorylated sites of YAP on S127 (**F**), was observed in P3 and P0+TGFβ1. Ratio of total YAP1 vs YAP1 phosphorylation plotted between YAP1/YAP1-S127 (**G**) showed no differences between groups upon fibroblast activation. Statistical significance was determined by one-way ANOVA with Tukey’s post-hoc test (n=3-4) *p ≤ 0.05, **p ≤ 0.01, ***p ≤ 0.001, ****p ≤ 0.0001, vs fibroblasts (P0).

## 4. Discussion

Our passaged hiPSC-CFs phenotypically resemble human or murine cardiac fibroblasts upon activation, which was evident in their key marker protein expression, such as TCF21, POSTN, and ED-A-Fn. Several studies have reported that primary human and murine cardiac fibroblasts attain a myofibroblast phenotype upon serial passaging, with reduced motility, incorporation of α-SMA into stress fibers, and formation of focal adhesions (Roche et al., 2016; Rohr, 2011; Rupert et al., 2020; Santiago et al., 2010). This is the first study to show that passaging hiPSC-derived cardiac fibroblasts similarly promotes the activation of fibroblasts and transition to a myofibroblast phenotype. We systematically captured snapshots of transcriptomic, proteomic, and metabolomic changes over three passages to understand the plasticity of hiPSC-CFs. This approach has provided insight into hiPSC-CFs and their use as ex vivo human models to study cardiac fibrosis and injury-related mechanistic aspects.

Cardiac fibroblasts undergo a surge in proliferation rate within 2-4 days of activation upon myocardial infarction or pressure overload injury, followed by a reduction in proliferation when they become myofibroblasts(Ali et al., 2014; Farbehi et al., 2019; Fu *et al*., 2018; Ivey et al., 2018; Ma et al., 2017; Moore-Morris et al., 2014). Several in vitro studies reported that while passaging murine primary adult cardiac fibroblasts, upon activation they undergo a proliferative burst followed by a significant reduction in the rate of proliferation when they attain the myofibroblast phenotype (Roche *et al*., 2016; Santiago *et al*., 2010). Similarly, in our hiPSC-CFs, we observed that proliferation indicators such as PCNA, MCM3, CDK1, and CDK4 were downregulated when passaged three times (P3), whereas TGFβ1-treated P0 fibroblasts displayed no significant difference in PCNA expression compared to P0. Our results suggest that passaging of hiPSC-CFs reduces their proliferative potential as they become more secretory and nonmotile myofibroblasts.

In the healthy myocardium, the ECM framework protects fibroblasts from stress. Injury or stress-related changes affect the ECM microenvironment and architectural integrity, and these changes promote fibroblast activation to become myofibroblasts (Burgess et al., 2002; Hinz and Gabbiani, 2003; Kural and Billiar, 2016; Snider et al., 2009; Wang et al., 2003). Fibroblast activation comes with notable changes in the secretion of ECM proteins compared to quiescent fibroblasts, which is reflected in altered gene expression of ED-A-Fn, Postn, Colα1, Colα3, integrins, and α-SMA in myofibroblasts (Davis and Molkentin, 2014; Ignotz and Massagué, 1986; Ivey and Tallquist, 2016; Serini et al., 1998; Snider *et al*., 2009; Tomasek et al., 2002). Lysyl oxidase and its family members are involved in the crosslinking of collagen and significantly contributes to cardiac remodeling(González-Santamaría et al., 2016). MMPs are also known to contribute to cardiac remodeling and the degradation of collagen during wound healing (DeLeon-Pennell et al., 2017; Visse and Nagase, 2003). In our hiPSC-CFs, similar changes were observed with passage-mediated fibroblast activation in Col1α1, Col1α2, Col4α1, ED-A-Fn, Postn, ITGA1, ITGB3, Lox, PLOD2, MMP-1, and MMP-2 expression, whereas Col3α1 and Vim expression were downregulated. Surprisingly, α-SMA mRNA expression was increased in P1 and P0+TGFβ1, but not in P2 or P3, whereas, α-SMA protein expression was elevated in P3, but not in P0+ TGFβ1. Indicating that α-SMA/ACTA2 showed a biphasic wave/effect in our mRNA and protein expression profiles at the time of sampling. However, phalloidin staining showed increased stress fiber incorporation in passage-mediated fibroblast activation. These results closely resemble the in vivo gene expression changes observed upon fibroblast activation to become myofibroblasts in the fibrotic myocardium.

Mitochondrial and metabolomic changes are crucial in determining the nature of fibroblast activation, persistence, and function in failing human hearts(Gibb *et al*., 2022). Fibrosis-associated myofibroblast transition is highly dependent on oxidative phosphorylation and increased mitochondrial content for energy production (Gibb et al., 2020; Negmadjanov *et al*., 2015). Glutaminolysis ensures the bioavailability of α-keto-glutarate (α-KG) to feed into the TCA cycle to increase collagen synthesis required for the ECM remodeling (Gibb *et al*., 2020; Lombardi et al., 2019). Myofibroblasts exhibit increased aerobic glycolysis and lactate production(Lombardi *et al*., 2019). During idiopathic pulmonary fibrosis, lung myofibroblasts enhance the glutaminolysis-mediated α-KG pathway for energy production that ensures the stability of collagen when remodeling occurs(Ge et al., 2018). Glutaminolysis inhibitors (e.g. CB-839 and BPTES, which inhibit GLS1) attenuate this process and collagen synthesis (Ge *et al*., 2018; Gibb *et al*., 2022). Our results provide evidence of similar metabolic changes during passage-mediated hiPSC-CFs activation to myofibroblasts, and notably, these changes were fully reversible by the GLS1 inhibitor BPTES.

Recent studies showed that in a murine MI model, increased collagen synthesis is correlated with higher glycolytic protein synthesis; this process is attenuated with the glycolysis inhibitor 2-deoxy-D-glucose (2-DG) which limited the activation of cardiac fibroblasts (Chen et al., 2021). In primary adult rat cardiac fibroblasts, passaging or TGFβ1 treatment promoted fibroblast activation and GLS1 expression; this process is limited by treatment with CB-839 (Chattopadhyaya *et al*., 2022). In our hiPSC-CFs upon passaging (P0 to P3) or TGFβ1 treatment, a gradual increase in mitochondrial oxygen consumption rate and extracellular acidification rate (glycolysis) were observed, indicating that quiescent cardiac fibroblasts are activated and utilizing high energy while they become myofibroblasts. However, BPTES, a glutaminolysis inhibitor, limited passage-mediated increases in OCR. Thus, activated hiPSC-CFs phenotypically resemble cardiac fibrosis-associated human or murine myofibroblasts.

TGFβ1 is a potent inducer of cardiac fibroblast to myofibroblast phenotype switching during pathological remodeling via canonical (Algeciras *et al*., 2021; Dobaczewski *et al*., 2010; Saadat *et al*., 2020; Villalobos *et al*., 2019) and non-canonical signaling (Zeglinski *et al*., 2016). Cardiac fibroblasts secrete the active form of TGFβ1 to promote myofibroblast activation (Sarrazy *et al*., 2014). TGFβ1I1 is also known as hydrogen peroxide inducible clone-5 (Hic-5), which was colocalized with α-SMA in human hypertrophic scars. In human dermal fibroblasts, mechanosensitive Hic-5 expression is dependent on TGFβ1 signaling and it was mediated by canonical Smad-3 and non-canonical MRTF-A and SRF pathways. Hic-5 plays a crucial role in the activation of myofibroblasts, stress fiber growth and assembly, and also nuclear translocation of MRTF-A (Dabiri et al., 2008; Varney et al., 2016). In our hiPSC-CFs, upon passaging, there was an induction of TGFβ1 and TGFb1I1/Hic-5 expression in myofibroblasts compared to non-passaged fibroblasts. These results are congruent with activation of the canonical TGFβ1-Smad signaling pathway, which potently promotes activation of fibroblasts and stress fiber assembly in cardiac myofibroblasts upon stress or injury.

Hippo signaling is also an important mechanism that activates fibroblasts to become myofibroblasts(Landry *et al*., 2021). YAP and TAZ bind together to translocate into the nucleus to activate the profibrotic signaling pathway, and phosphorylation of YAP hinders nuclear translocation in fibroblasts and myofibroblasts. We found that passage-mediated fibroblast activation in hiPSC-CFs resulted in increased TAZ expression and dephosphorylation of YAP1 at S127, which may induce YAP1 nuclear translocation along with TAZ to promote profibrotic gene expression.

To mimic fibrosis-associated fibroblast activation and other cellular changes in cardiac fibroblasts, in vitro experiments have been largely focused on murine primary fibroblasts (Gilles *et al*., 2020). Most commercially available human primary cardiac fibroblasts (HCFs) are very limited in cell numbers and are passaged more than once prior to freezing. Frozen cells need to be further expanded to increase the cell numbers before initiating the experiments. Fibroblasts are highly mechano-sensitive in nature, and upon passaging are activated and transition towards the myofibroblast phenotype which was clearly evident in our hiPSC-CFs. The main advantages over murine fibroblasts are avoidance of batch-to-batch variability and low passage number, whereas hiPSC-CFs can be maintained up to five passages quiescently using the TGFβ1inhibitor SB431542 (Zhang *et al*., 2019a). In our hands, hiPSC-CFs can be obtained in larger quantities, which helps with conducting multiple experiments and with reproducibility of results, as our cells exhibited relatively low variability across experiments. hiPSC-CFs can be cryopreserved at the P0 stage, which will help avoid batch variations. Mimicking the profibrotic condition with proper controls in HCFs is very unlikely due to limited cell numbers as well as nature of the cells after passaging.

A major advantage of hiPSC-derived cells is that they can be bioprinted as three-dimensional cardiac tissue-like structures, consisting of cardiomyocytes, fibroblasts, and endothelial cells together from the same patient-specific hiPSC cells, and can mimic the cardiac fibrotic condition inside the myocardium. These patient-specific hiPSC-derived cells will help to better understand gene related abnormalities and the severity of disease phenotypes with detailed signaling mechanisms for potential therapeutic applications. Thus hiPSC-CFs offer distinct advantages over murine or human cardiac fibroblasts.

The present study shows that passage-mediated hiPSC-CFs activation aligns well with the in vivo physiology of fibrotic myocardium, as shown by transcriptomic, proteomic and metabolomic analysis. Mimicking profibrotic conditions in hiPSC-CFs is highly feasible as shown by our analyses, and more physiologically relevant to humans than murine primary fibroblasts. Most importantly, in our hiPSC-CFs, in P0+TGFβ1, TGFβ1 activates the fibroblasts and their transcriptomic, proteomic and metabolomic profile shows that they are in the continuum phase to become myofibroblasts but are not identical to P3 fibroblasts. Our study results will help clarify the misrepresentations of phenotype and disparity in previous studies.

## 5. Experimental procedures

Resource availability

### Lead contact

Request for resources, reagents, and protocols should be addressed to the corresponding author, GF Tibbits, PhD.

### Materials availability

This study did not generate new unique reagents.

### Data and code availability

All data presented in this study are available in the main text, and from the corresponding author upon request.

### Human induced pluripotent stem cell maintenance and fibroblast differentiation

We used two hiPSC lines (iPS IMR90-1 and STAN248i-617C1), obtained from the Wicell Research Institute, for fibroblast differentiation, which was accomplished using a few modifications of a previously published protocols (Zhang *et al*., 2022; Zhang *et al*., 2019a). The hiPSCs were maintained on Corning Matrigel-coated 6-well tissue culture plates with mTeSR-Plus medium (STEMCELL Technologies). Cells were passaged every 4 days using ReLeSR media (StemCell Technologies) and were then seeded on a six-well Matrigel-coated plate at a density of 175,000 cells cm^−2^ in mTeSR-Plus medium. For fibroblast differentiation, on day-0 >90% confluent hiPSC monolayers were treated with 6 μM CHIR99021 containing RPMI-1640+B27-insulin media for 48 hrs. On day-2 the media was replaced with RPMI-1640+B27-insulin media for 24 hrs; on day-3, cells were treated with 5 μM of the WNT signaling inhibitor IWR1 (I0161, Sigma) containing RPMI-1640+B27-insulin media for 48 hrs. On day-5, hiPSC-derived progenitor cells (hiPSC-PCs) were passaged and replated at a density of 100,000 cells cm-2 in advanced DMEM medium (12634028, Gibco) consisting of 1% FBS and gluMax along with 5 μM CHIR99021 and 2 μM retinoic acid (R2625, Sigma-Aldrich) for 72 hrs. On day-9, cells were maintained in advanced DMEM consisting of 1% FBS and gluMax for 48 hrs. On day-11, hiPSC-derived epicardial cells (hiPSC-EPCs) were passaged and replated at a density of 100,000 cells cm-2 then treated with 10 ng/ml FGF2 (100-18B, PeproTech) and 10 μM SB431542 (S1067, Selleck chemicals) in fibroblast growth medium-3 (FGM-3) (PromoCell) for another 6 days, with the media changed every 48 hrs. On day-20, hiPSC derived fibroblasts (hiPSC-FBs) (P0) were ready to be passaged and replated at a density of 75,000 cells cm-2 with FGM-3 media. Then after every 96 hours hiPSC-FBs were passaged until passage-3 (P3). We considered each differentiation into a replicate and multiple differentiations were carried out with two different hiPSC lines. For the TGFβ1 treatment, hiPSC-FBs were serum starved for 6 hrs without serum supplement in FGM-3 media, prior to the treatment with 10 ng/ml of TGFβ1 for 48 hrs.

### Protein isolation and purification

The hiPSC-Fb proteins were isolated from cell pellets using TrypLE dissociation reagent, which was added to each well and incubated for ∼5-6 minutes. This was followed by media neutralization and the collection of detached cells. The homogenous cell suspension was centrifuged to obtain the cell pellet. The pellet was then resuspended in an aliquot of lysis buffer in which the volume was proportional to the number of cells. The cells were counted using the CellDrop automated cell counter. The lysis buffer contained 100 mM HEPES (pH= 8-8.5) + 20% SDS. Cell lysates were then placed for 5 minutes in 95°C water bath, followed by 3 minutes on ice. To complete the lysis, the samples were sonicated twice for 30 seconds, and placed on ice between rounds. This was followed by treatment with benzonase (EMD Millipore-Sigma) at 37°C for 30 min to fragment the chromatin.

After obtaining the lysates, each sample was reduced with 10 mM dithiothreitol (DTT) at 37°C for 30 min, followed by alkylation with 50 mM chloroacetamide (CAA) for 30 min in the dark. The alkylation was quenched in 50 mM DTT for 10 min at room temperature. To each sample, hydrophilic and hydrophobic Sera-Mag Speed Beads (GE Life Sciences, Washington, US) were added in 1:1 ratio. Proteins were then bound to the beads with 100% ethanol (80% v/v) and washed twice with 90% ethanol.

After binding of the proteins to the beads, the samples underwent an overnight enzymatic digestion with trypsin (1:50 w/w), followed by C18 matrix midi-column clean up and elution. The BioPure midi columns (Nest Group Inc.) were conditioned with 200 μL methanol, 200 μL 0.1% formic acid (FA), 60% acetonitrile (ACN), and 200 μL 0.1% trifluoroacetic acid (TFA). The sample pH was adjusted to pH=3 – 4 using 10% TFA prior to loading the columns. Samples were sequentially eluted 3 times with 70 μL, 70 μL, and 50 μL 0.1% FA, 60% ACN. The eluate was collected into LoBind tubes and placed into a SpeedVac to remove the organic ACN. Lastly, the samples were resuspended in 0.1% FA (ready for MS injection).

### Protein detection and quantification using LC/MS

The total protein concentration for each of the purified samples was determined using Nanodrop. Mass spectrometric analyses were performed on Q Exactive HF Orbitrap mass spectrometer coupled with an Easy-nLC liquid chromatography system (Thermo Scientific). The time of flight was calculated as the period between injection to detection. In the MS sample chamber (96-well plate), the samples were completely randomized across all conditions and biological replicates. A total of 1 μg peptides per sample was injected for analysis. The peptides were separated over a three-hour gradient consisting of Buffer A (0.1% FA in 2% ACN) and 2%-80% Buffer B (0.1% FA in 95% ACN) at a set flow rate of 300 nL/min. The raw mass-to-charge (M/Z) data acquired from the Q Exactive HF were searched with MaxQuant (MQ)-version 2.0.0.0 (Max Planck Institute, Germany), and Proteome discoverer software (Thermo Fisher Scientific, CA, USA), using the built-in search engine, and embedded standard DDA settings. The false discovery rate for protein and peptide searches was set at 1%. Digestion settings were set to trypsin. Oxidation (M) and Acetyl (N-term) were set as dynamic modifications. Carbamidomethyl (C) was set as fixed modification, and Phosphorylation (STY) was set as a variable modification. For the main comparative protein search, the human proteome database (FASTA) was downloaded from Uniprot (2022_06; 20,365 sequences) and used as the reference file. Common contaminants were embedded from MaxQuant. Peptide sequences that were labeled as "Potential contaminant/REV" were excluded from the final analysis.

### Seahorse assay

hiPSCs were differentiated 4 days apart to attain all the passages (P0, P1, P2 and P3) on the same day to perform Seahorse assay. hiPSC-derived fibroblasts were dissociated using TrypLE and replated at a density of 15,000 cells/well on Matrigel-coated Seahorse Xfe96 well cell culture plates. Cells were serum starved for 6 hrs without serum supplement in FGM-3 media, after 24 hrs of initial plating, followed by treatment with 10 ng/ml TGFβ1 for 48 hrs or 10 μM BPTES treated after 24 hrs of TGFβ1 treatment for 24 hrs. On the day of the experiment, the cell medium was changed to Seahorse media containing RPMI1640 medium without glucose and sodium bicarbonate (R1383, Sigma) supplemented with glucose (10 mM, Sigma), glutamine (2 mM, Sigma), and sodium pyruvate (1 mM, Sigma) and cells were incubated in a non-CO_2_ incubator for a maximum of 45 minutes. Subsequently, the Seahorse Mito Stress test was performed using Agilent Seahorse XFe96 Analyzer by sequential injection of ATP synthase inhibitor oligomycin (1 μM, Sigma), mitochondrial uncoupler FCCP (1.5 μM, Sigma), and mitochondrial complex I and III inhibitors rotenone (100 nM, Sigma)/antimycin A (1 μM, Sigma). The raw Oxygen Consumption Rate (OCR) and Extra Cellular Acidification Rate (ECAR) values were normalized based on cell numbers per each well using Wave software (Agilent Technologies, Inc). Data analysis and visualization were performed using GraphPad Prism (Version 9.5.1 (528)).

### Immunofluorescence staining

hiPSC-derived fibroblasts were cultured on a cover glass coated with Matrigel with appropriate seeding density. Cells were washed thrice with PBS followed by 4% PFA for 10 mins at room temperature (RT). Fixed cells were quenched with 100 mM glycine for 5 mins at RT, then washed with PBS thrice. Cells were permeabilized using 0.1-0.3% triton X-100 in PBS for 3 mins, followed by washing with PBS thrice. Permeabilized cells were then blocked with 5% BSA for 45 mins, followed by overnight incubation with 1:500 dilution of DDR-2 antibody (ab63337, Abcam). The cover glass with the cells was then washed with PBS 3 times for 5 mins in an orbital motion shaker at 50-60 rpm. The cells were incubated with phalloidin (A34055, ThermoFisher) and 1:500 dilution of goat anti-rabbit secondary antibody in 1% BSA for an hour at RT in the dark. The cells were washed with PBS for 5 mins three times in an orbital motion shaker, then mounted on a glass slide using ProLong Glass Antifade Mountant (P36980, ThermoFisher) and dried for 2 hours, then imaged under a Nikon Ti microscope.

### Quantitative real-time PCR

hiPSC-derived fibroblasts were dissociated using TrypLE, and the cells were pelleted by centrifugation and flash-frozen prior to RNA isolation. Total RNA from flash-frozen cells was processed using NucleoMag (744350.1, Takara) Magnetic bead-based RNA isolation following the manufacturer’s instructions. cDNA was generated from 1 µg RNA samples using the iScript cDNA Synthesis kit (Bio-Rad, USA). qPCR reactions were prepared using 5 ml SsoAdvanced Universal SYBR Green Supermix (Bio-Rad, USA) and 4 ml 1:6 diluted cDNA template along with 200 nM forward and reverse primers in a total volume of 10 ml per reaction. PCR amplification was performed in duplicate for each reaction on a CFX384 Touch Real-Time PCR (Bio-Rad, USA). The cycling conditions were 95°C (3 min), followed by 40 cycles of denaturation at 95°C (15 s) and extension at 62°C (30 s). After amplification, a continuous melt curve was generated from 60 to 95°C to confirm the amplification of single amplicons. Relative gene expression was calculated using the 2−ΔΔCt method with normalization to GAPDH (primer sequences are in Supplemental Table 1).

### NanoString analysis for mRNA expression profile

Total RNA from flash-frozen cells was processed using NucleoMag (744350.1, Takara) Magnetic bead-based RNA isolation, according to the manufacturer’s instructions. The concentration and purity of the RNA were determined using a NanoDrop ND-1000 spectrophotometer (Thermo Fisher Scientific). Multiplexed mRNA profiling was conducted using a human fibrosis V2 panel codeset containing 762 fibrosis-specific gene probes synthesized by NanoString Technologies Inc. and read on the NanoString nCounter® SPRINT Profiler. A total of 50 ng purified RNA per sample was hybridized overnight (16 h) to the custom capture and reporter probes. Hybridized samples were loaded into each channel of the nCounter® SPRINT cartridge. Raw mRNA counts were collected, and the results were normalized to ten housekeeping genes (ACAD9, ARMH3, CNOT10, GUSB, MTMR14, NOL7, NUBP1, PGK1, PPIA, RPLP0). Analysis was performed on the nSolver analysis software and the Advanced Analysis module (NanoString Technologies Inc.).

### Statistical analysis

Data are reported as mean ± standard deviation of a minimum of three independent biological replicates. To reduce variability, all cells, including control and treatment groups, were isolated and cultured on the same day. For data analysis, different treatments and samples between groups were kept blinded. Violin plots from NanoString and mass spectrometry data sets were generated and analyzed using GraphPad prism (9.5.1) software with one-way analysis of variance with Tukey’s post-hoc analysis as appropriate, with P<0.05 considered to be statistically significant. Principle component analysis (PCA) was performed based on the Benjamini-Hochberg method false discovery rate (FDR) adjusted to P-value < 0.05. Group correlation heatmap was based on Pearson correlation and FDR adjusted to P-value < 0.05. Volcano plots were prepared based on Welch’s t-test and adjusted P < 0.05 using Perseus v2.0.10.0 software. Heatmap for selected genes was generated with the pheatmap package (clustering_distance_rows = "euclidean") and with a P value of <0.05 considered statistically significant. The compareCluster function was used to compare the top 10 gene ontology (GO) terms between each group (pAdjustMethod = "BH", pvalueCutoff = 0.05, qvalueCutoff = 0.2). Gene set enrichment analysis was performed using the gseGO function (pvalueCutoff = 0.05). Dot s were generated using ggplot2 dotplot (Wickham, 2016).

## Acknowledgments

GFT research is supported by grants from the Canadian Institutes of Health Research (CIHR PJT-488595) and the Stem Cell Network (FY21 / ACCT2-13). The RAR research program is supported by The Heart and Stroke Foundation of Canada (G-22-0032033) and CIHR grants (PJT166105 and PJT180474). RSN is supported by postdoctoral research fellowships from CIHR and Michael Smith Health Research BC. Most importantly, we appreciate the incredible infrastructural support from BC Children’s Hospital Foundation and the Mining for Miracles (MoM) for the creation of the Cellular and Regenerative Medicine Centre (CRMC) at the BC Children’s Hospital Research Institute.

## Author contributions

R.S.N., optimized the culture conditions for cardiac fibroblast differentiation and developed the methodology, performed experiments, and wrote the manuscript under the supervision of G.F.T. F.J., H.H., and D.H.B., assisted with experiments. F.J., H.H., S.D., and C.L., helped in data analysis. I.M.C.D., R.A.R., M.P.C., and G.F.T. revised the manuscript.

