## Supplemental file for "Molecular and metabolomic characterization of hiPSC-derived cardiac fibroblasts transitioning to myofibroblasts"

| | |
|---|---|
| ED-A-Fn-F (human) | 5’- ACTGCAGTAACCAACATTGATC -3’ |
| ED-A-Fn-R (human) | 5’- CACCCTGTACCTGGAAACTTGC -3’ |
| Postn-F (human) | 5’- TGTTGCCCTGGTTATATGAG -3’ |
| Postn-R (human) | 5’- GTGGTGGCTCCCACGATGCC -3’ |
| TCF21-F (human) | 5’- CACTTGAGGCAGATCCTGGCTA -3’ |
| TCF21-R (human) | 5’- CGGTCACCACTTCTTTCAGGTC -3’ |
| β-actin-F (human) | 5’- ATTGCCGACAGGATGCAGAA -3’ |
| β-actin -R (human) | 5’- GGGCCGGACTCGTCATACTC -3’ |

**Supplemental Table 1**. Primers used for quantitative real-time PCR. Forward (F) and reverse (R) primers are shown.

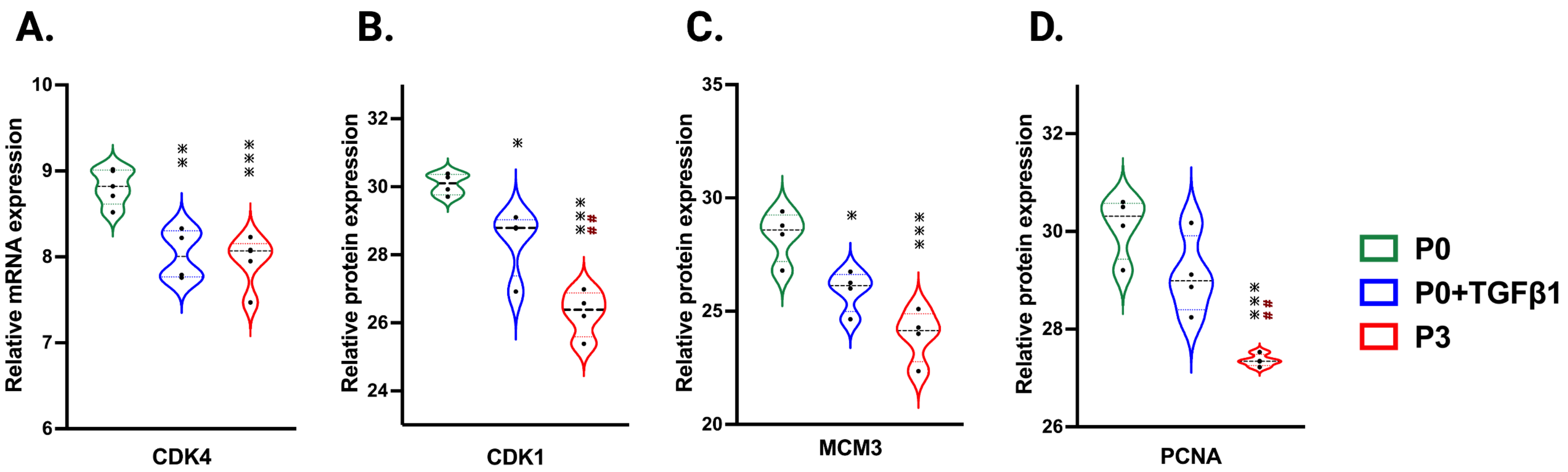


**Figure S-1: Passaging influences markers of fibroblast proliferation**. Violin plots represent mRNA expression of CDK4 (A), and proteomic expression of PCNA (B) MCM3 (C) and CDK1 (D), showed reduction in the proliferation in P3 and P0+TGFβ1 compared to P0. Statistical significance was determined by one-way ANOVA with Tukey's post-hoc test (n=4-5) *$p \leq 0.05$, **$p \leq 0.01$, ***$p \leq 0.001$, ****$p \leq 0.0001$, vs fibroblasts (P0) and $^{\#\#}p \leq 0.01$, vs P0+TGFβ1.

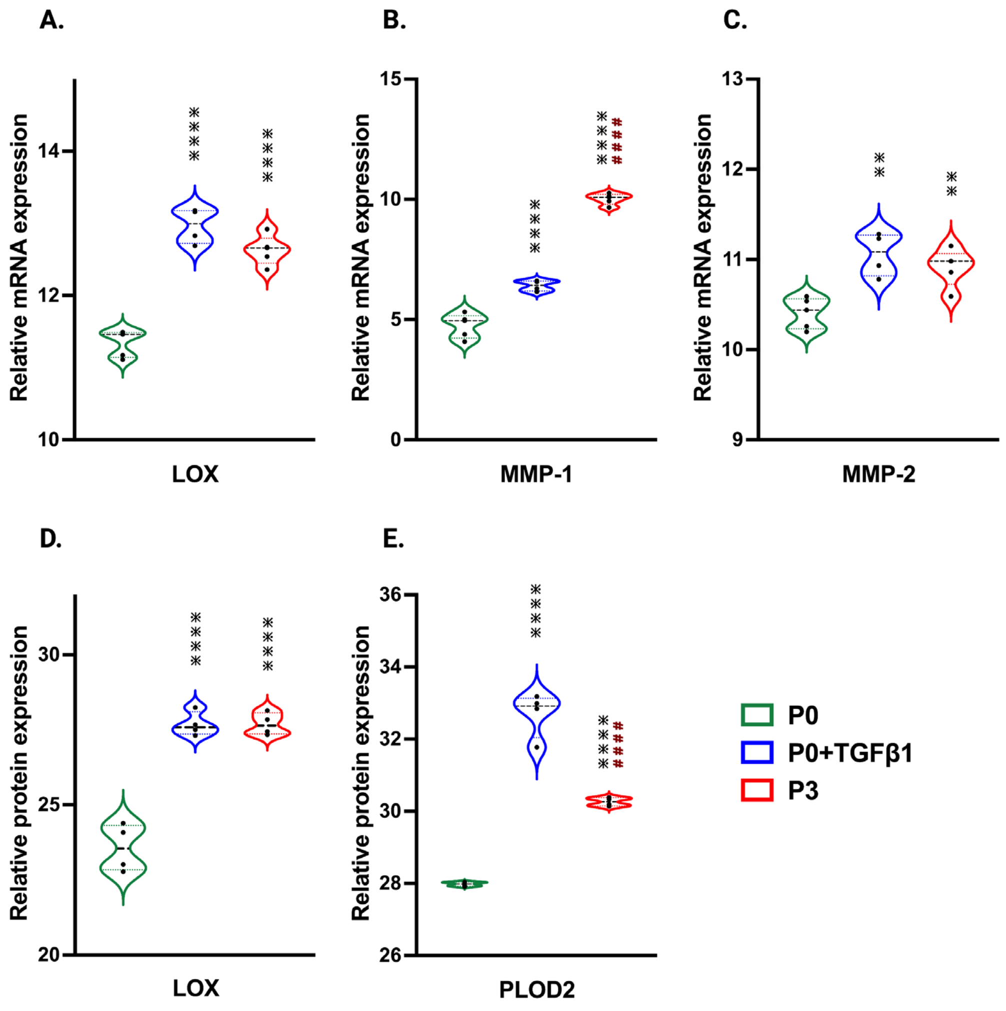


**Figure S-2: Passaging augments ECM remodeling enzyme expression**. Violin plots represent mRNA expression of LOX (A), MMP-2 (B) MMP-1(C) and proteomic expression of LOX (D), and PLOD2 (E), showing elevated ECM remodeling enzymes as an indirect measure of collagen synthesis in P3 and P0+TGFβ1 compared to P0. Statistical significance was determined by one-way ANOVA with Tukey's post-hoc test (n=4-5) *$p \le 0.05$, **$p \le 0.01$, ***$p \le 0.001$, ****$p \le 0.0001$, vs fibroblasts (P0) and $^{####}p \le 0.0001$ vs P0+TGFβ1.

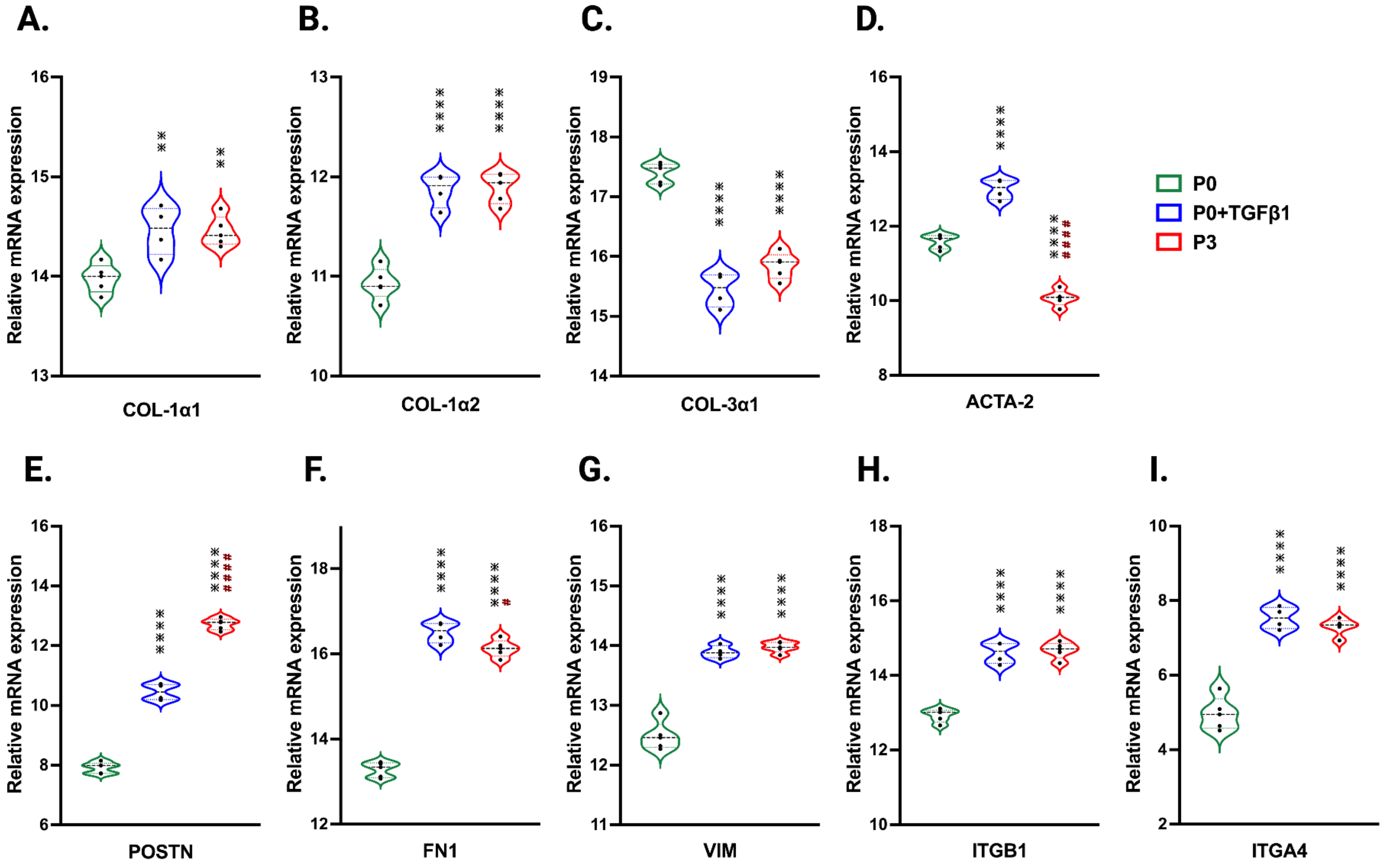


**Figure S-3: Passaging promotes profibrotic gene expression in myofibroblasts**. Violin plots of profibrotic gene expression using Nanostring analysis in P0+TGFβ1 and P3 compared to P0, (A-I); Statistical significance was determined by one-way ANOVA with Tukey's post-hoc test (n=4-5). $*p<0.05$, $**p<0.01$, $***p<0.001$, $****p<0.0001$ vs fibroblasts (P0) and $^{\#}p \leq 0.05$, $^{\#\#}p \leq 0.01$, $^{\#\#\#\#}p \leq 0.0001$ vs P0+TGFβ1.

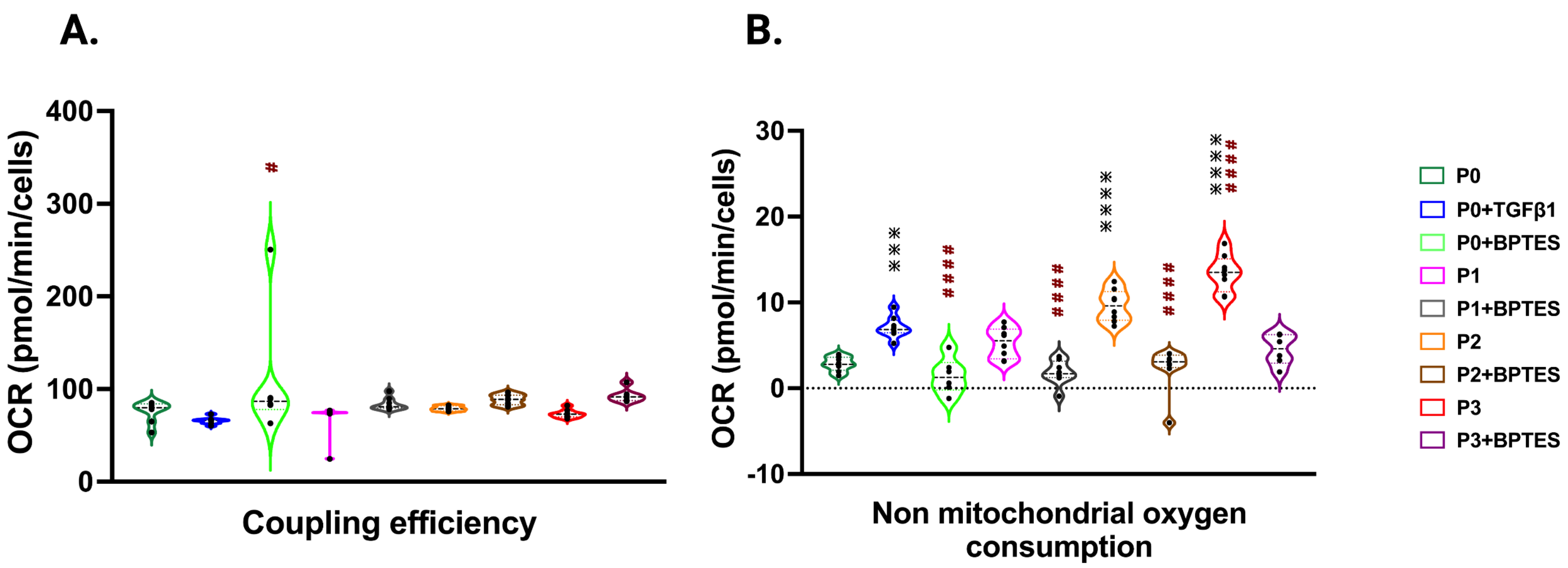


**Figure S-4**: **Passage-induced fibroblast activation and mitochondrial metabolism**. Seahorse assay reveals that coupling efficiency (A) showed no differences regardless of passage or treatment also, non-mitochondrial OCR (B) varied with passage (P3) and/or TGFβ1 treatment, whereas the glutaminase inhibitor BPTES inhibited this process when compared to non-passaged fibroblasts (P0). Statistical significance was determined by one-way ANOVA with Tukey's post-hoc test (n = 3-8); *p ≤ 0.05, **p ≤ 0.01, ***p ≤ 0.001, ****p ≤ 0.0001, vs fibroblasts (P0) and $^{\#}$p ≤ 0.05, $^{\#\#\#\#}$p ≤ 0.0001 vs P0+TGFβ1.
